# Function of the alternative electron transport chain in the *Cryptosporidium parvum* mitosome

**DOI:** 10.1101/2024.10.01.616074

**Authors:** Silu Deng, L. David Sibley

## Abstract

*Cryptosporidium parvum and C. hominis* possess a remanent mitochondrion called the mitosome, which lacks DNA, the tricarboxylic acid cycle, a conventional electron transport chain, and ATP synthesis. The mitosome retains ubiquinone and iron sulfur cluster biosynthesis pathways, both of which require protein import that relies on the membrane potential. It was previously proposed that the membrane potential is generated by electrons transferred through an alternative respiratory pathway coupled to a transhydrogenase (TH) that pumps hydrogens out of the mitosome. This pathway relies on an alternative oxidase (AOX) and type II NADH dehydrogenase (NDH2), which also exists in plants, some fungi, and several protozoan parasites. To examine this model, we determined the location and function of AOX and NDH2 in *C. parvum*. Surprisingly, we observed that NDH2 was localized to parasite surface membranes instead of the mitosome. Furthermore, a Δ*ndh2* knockout (KO) strain was readily obtained, indicating that this protein is not essential for parasite growth. Although, AOX exhibited a mitosome-like staining pattern, we readily obtained an Δ*aox* knockout strain, indicating that AOX is also dispensable for parasite growth. The growth of the Δ*aox* strain was inhibited by the AOX inhibitors SHAM and 8-HQ to the same extent as wild type, indicating that AOX is not the target of these inhibitors in *C. parvum*. Collectively, our studies indicate that NDH2 and AOX are non-essential genes in *C. parvum*, necessitating an alternative mechanism for maintaining the mitosome membrane potential.

**Importance:** Cryptosporidiosis is the leading cause of diarrhea in young children and immunocompromised individuals, particularly AIDS/HIV patients. The only FDA approved drug against cryptosporidiosis, nitazoxanide, has limited effectivity in immunocompromised patients and is not approved for usage in children under 1 year old. Genomic analysis and previous studies proposed an alternative respiration pathway involving alternative oxidase (AOX) and type II NAD(P)H dehydrogenase (NDH2), which are thought to generate the mitosome membrane potential in *C. parvum*. Additionally, AOX and NDH2 were nominated as potential drug targets, based on their absence in mammalian hosts and sensitivity of parasite growth to known inhibitors of AOX. However, our study demonstrated that NDH2 is not localized in mitosome, AOX non-essential for parasite growth, and knockout lines lacking this enzyme are equally sensitive to AOX inhibitors. These findings indicate that AOX and NDH2 are not ideal candidates for future drug development against cryptosporidiosis and force a re-evaluation for models of how the mitosome generate its membrane potential.

## Introduction

Mitochondria originated from α-proteobacterial endosymbionts and serve as ATP-generating factories in aerobic eukaryotes (1). During evolution, their genomes and proteomes have radically evolved, affecting biological processes and structure. Despite the trend toward increased genome complexity, many parasitic and symbiotic organisms have reduced mitochondrion-related organelles (2, 3). The reduction primarily manifests in the loss of mitochondrial genome, the simplified structure, decreased size, and specialized or diminished functionality, which often makes it difficult to recognize and identify these relict mitochondrial compartments. The function of mitochondrion-related organelles ranges widely across different organisms, including hydrogenosomes comprising anaerobic energy metabolism and the mitosomes retaining the Fe-S biosynthesis with a loss of energy generating machinery (4).

In apicomplexan parasites, the morphology and functionality of mitochondria vary from organism to organism, species to species, and in different life cycle stages. The *Plasmodium falciparum* mitochondrion appears as a single, small and discrete organelle during the ring and early trophozoites stages, but undergoes branching and elongation in the transition between mature trophozoite and schizont (5). The mitochondrion lies close to the apicoplast throughout the entire asexual life cycle (5). The mitochondria of *Toxoplasma gondii* undergoes a reversible change from a collapsed tubular structure to peripheral and lasso-shaped organelle during the transition from extracellular to intracellular stages (6, 7). Mitochondria in both *Plasmodium* spp. and *T. gondii* maintain an electron transport chain (ETC) and integrated tricarboxylic acid (TCA) cycle for ATP synthesis that is similar to that of most other eukaryotes with the presence of an alternative complex I, and type II NADH dehydrogenase (NDH2), replacing the canonical complex I in the ETC (8, 9). In contrast, the genus *Cryptosporidium* possesses mitochondrial relicts that are much smaller in size and have reduced functionality (10, 11). The gastric-dwelling *Cryptosporidium* species, including *C. muris* and *C. andersonii*, contain a functional ETC and complete TCA cycle for energy metabolism. In contrast, the intestine-dwelling species, particularly *C. parvum* and *C. hominis*, lack DNA, the TCA cycle and retain only two subunits of the ATPase, and are thus incapable of generating ATP. Instead, they have an alternative ETC consisting of two dehydrogenases, NDH2 and malate-quinone oxidoreductase (MQO), as well as an alternative oxidase (AOX) (4, 11). This remnant organelle is referred to as the mitosome, which is a roughly spherical, 150-300 nm in diameter, double membrane-bounded organelle between the nucleus and crystalloid body at the posterior end of the sporozoites (12).

NDH2 proteins have been widely described in plants, fungi, and protists, but are absent in mammalian mitochondria (13). Many plants, fungi and protozoa possess both complex I and NDH2, while apicomplexan parasites only retain NDH2 (14). In the modified ETC, NDH2 catalyzes the oxidation of NADH to NAD^+^, followed by reduction of quinone to quinol, resulting in electron transfer from NADH to quinone (15). However, unlike the canonic complex I, NDH2 is embedded in the inner leaflet of the inner mitochondrial membrane rather than being a transmembrane protein, and thus it cannot directly mediate proton-pumping (15, 16). Due to their absence in mammals, type II NADH dehydrogenases have been proposed as attractive drug targets against *Mycobacterium tuberculosis* (17) and as potential drug targets against apicomplexan parasites (18). MQO is another possible electron donor for the mitosome ETC encoded in *C. parvum*. MQO catalyzes the reversible NAD^+^-dependent oxidation of malate to oxaloacetate and mediates electron transport through the reduction of quinone in mitochondria, which has been reported to be a pH-dependent process in functional analysis in vitro (19, 20). The only electron acceptor identified from the genome of *C. parvum* is AOX, a cyanide-resistant ubiquinol oxidase found in the mitochondria of all the plants as well as some fungi and protozoa, which can accept electrons directly from coenzyme Q and catalyze the reduction of oxygen to water (21, 22). AOX is not expressed in either *T. gondii* or *Plasmodium* spp., while it plays a crucial role in respiration and development of both bloodstream and procyclic forms of *Trypanosoma brucei*, making it a viable chemotherapeutic target for African trypanosomiasis (23). A previous study has reported that the AOX inhibitors salicylhydroxamic acid (SHAM) and 8-hydroxyquinoline (8-HQ) inhibit the growth of *C. parvum* in cell culture (24, 25), leading to the suggestions that AOX could be a potential drug target for cryptosporidiosis. Membrane-bound NAD(P) transhydrogenase (TH) facilitates the reversible transfer of hydride ions between NAD(H) and NADP(H) while simultaneously translocating protons across the membrane, an activity that is conserved across prokaryotes and eukaryotes (26). In mammals, TH proteins reside in the mitochondria inner membrane (27), whereas *Plasmodium* TH is found in the apicoplast rather than mitochondria (28). It was previously proposed that the TH may be coupled to the alternative ETC in *Cryptosporidium*, thus generating the mitosome membrane potential (8).

Although several models have been put forward suggesting a role of NDH2 and AOX in generating the mitosome membrane potential based on genome comparisons and analogy to other organisms (4, 8, 11, 29, 30), no studies have investigated the localization and essentiality of these components. In present study, we localized the AOX and NDH2 proteins using epitope tags and tested their essentiality for growth. Our findings indicate that NDH2 is not localized in the mitosome and neither NDH2 nor AOX are essential for growth of *C. parvum*, forcing a revision of current models for how the mitosome membrane potential is generated.

## Results

### Proposed model for electron transport chain in mitosome of *C. parvum*

Genomic analysis of *C. parvum* and *C. hominis* revealed a progressive reduction in mitochondrial functions in these species that infect the small intestine relative to *C. muris* that resides in the stomach (11, 29, 30). Although *C. parvum* retains genes encoding the proteins involved in the ubiquinone biosynthesis, only a few enzymes mediating the ETC were identified in *C. parvum*, including MQO, NDH2, and AOX (4, 11, 30). Thus, a simplified model for an alternative electron transport chain in *C.parvum* mitosome was proposed in previous reviews based on comparative genomics (4, 8). In this model, electrons are produced during oxidation of malate to oxaloacetate by MQO, or dehydrogenation of NAD(P)H to NAD(P)^+^ by NDH2, and then transferred to CoQ, which release the electrons to AOX via the reduction of quinone to quinol (**Fig. 1A**). AOX subsequently catalyzes the oxidation of quinol and the reduction of oxygen to water and this alternative ETC is coupled to proton pumping by TH (**Fig. 1A**). Two MQO-like proteins (encoded by *cgd7_470* and *cgd7_480*), two TH-like proteins (encoded by *cgd1_990* and *cgd8_2330*) one NDH2 (encoded by *cgd7_1900*), and one AOX (encoded by *cgd3_3120*) are identified from the genomic analysis of *C. parvum* (11, 26). The HyperLOPIT proteomic database (31) indicated a nuclear/cytoplasmic location for both MOQs, microneme location for TH (*cgd8_2330*) and unassigned for TH (*cgd1_990*), while NDH2 was found with the inner membrane complex (IMC) and AOX fractionated with the mitosome, respectively (**Fig. 1B**). Here, we primarily focused on characterizing the localizations and functions of NDH2 and AOX in *C. parvum*.

**Figure 1.**
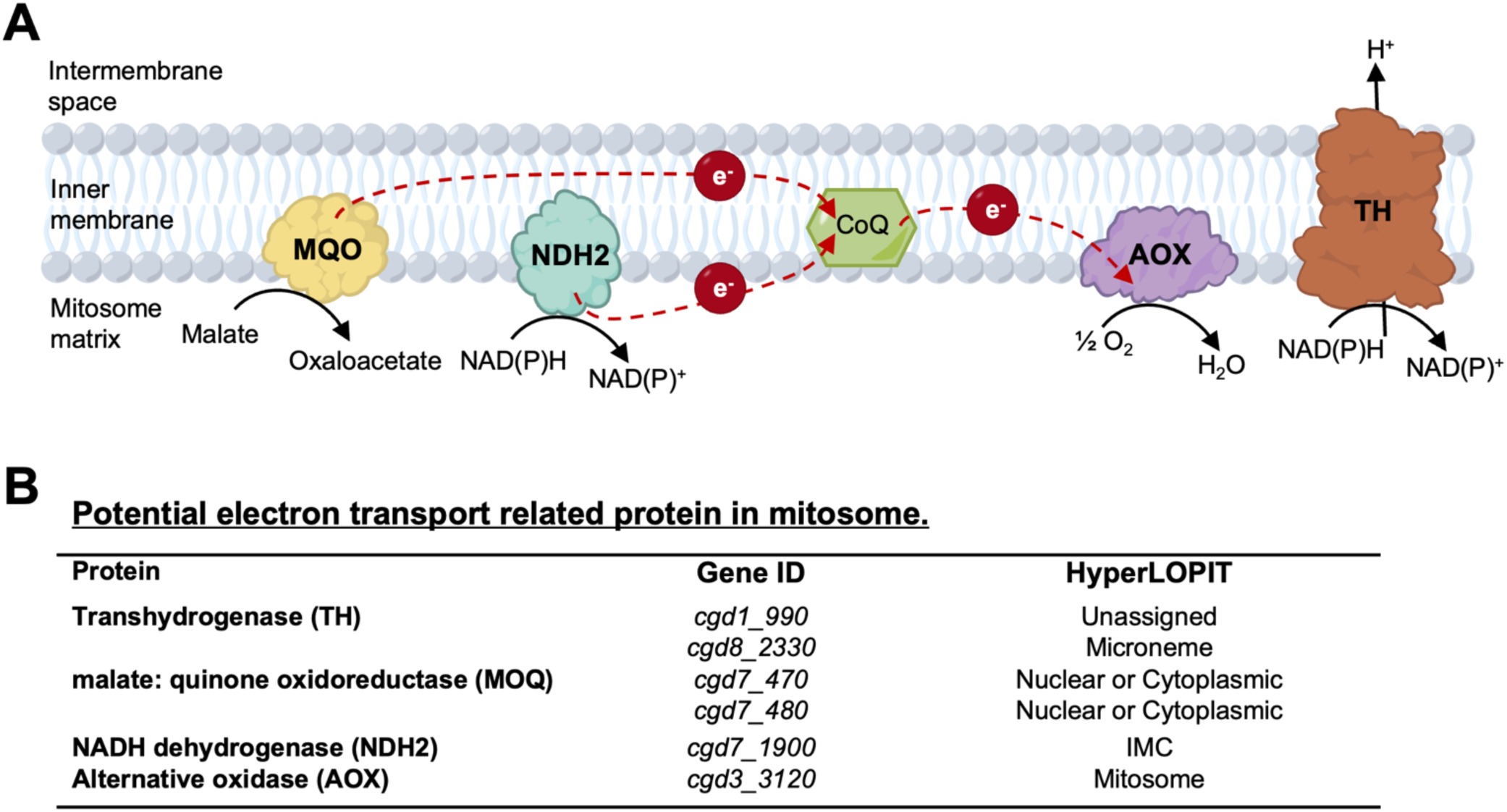
Proposed model for electron transport chain in *C. parvum* mitosome. (A) Schematic diagram of proposed model for electron transport chain in *C. parvum* mitosome. MQO, malate: quinone oxidoreductase; NDH2, type II NAD(P)H dehydrogenase; CoQ, coenzyme Q; AOX, alternative oxidase. (B) Summary of annotated genes from genomic and proteomic databases for *C. parvum.* Gene IDs were obtained from NCBI gene database (https://www.ncbi.nlm.nih.gov/ gene). Predicted localization was based on the HyperLOPIT proteomic dataset (31).

### CpNDH2 is present at the parasite surface

To investigate the localization of CpNDH2, we employed CRISPR/Cas9 gene editing to add a triple hemagglutinin (3HA) epitope tag to the C terminus (**Fig. 2A**). The tagging construct also contained a selection cassette consisting of nanoluciferase (Nluc) and neomycin resistance (Neo^R^) jointed by a split peptide motif (P2A) and driven by an enolase promoter (**Fig. 2A**). The CpNDH2-3HA tagging plasmid was co-transfected into excysted sporozoites with a CRISPR/Cas9 plasmid containing a gene-specific sgRNA targeting Cp*NDH2*. Transfected parasites were selected by puromycin in Ifng^-/-^ mice followed by the second round of selection and amplification in Nod scid gamma (NSG) mice. The genotype of transgenic parasites was validated using diagnostic PCR to detect insertion of the tag at the endogenous locus using with DNA extracted from mice fecal pellets at 30 days post infection (dpi) (**Fig. 2B**). Growth of the CpNDH2-3HA tagging strain was assessed by testing the luminescence signal from Nluc gene from NSG fecal pellets at from 3 to 30 dpi (**Fig. 2C**). To visualize the location of CpNDH2, we performed immunofluorescence assays (IFA) using the CpNDH2-3HA tagged parasite. CpNDH2 protein expression was detected in both trophozoite and meront stages, where it showed a surface membrane staining pattern (**Fig. 2D**).

**Figure 2.**
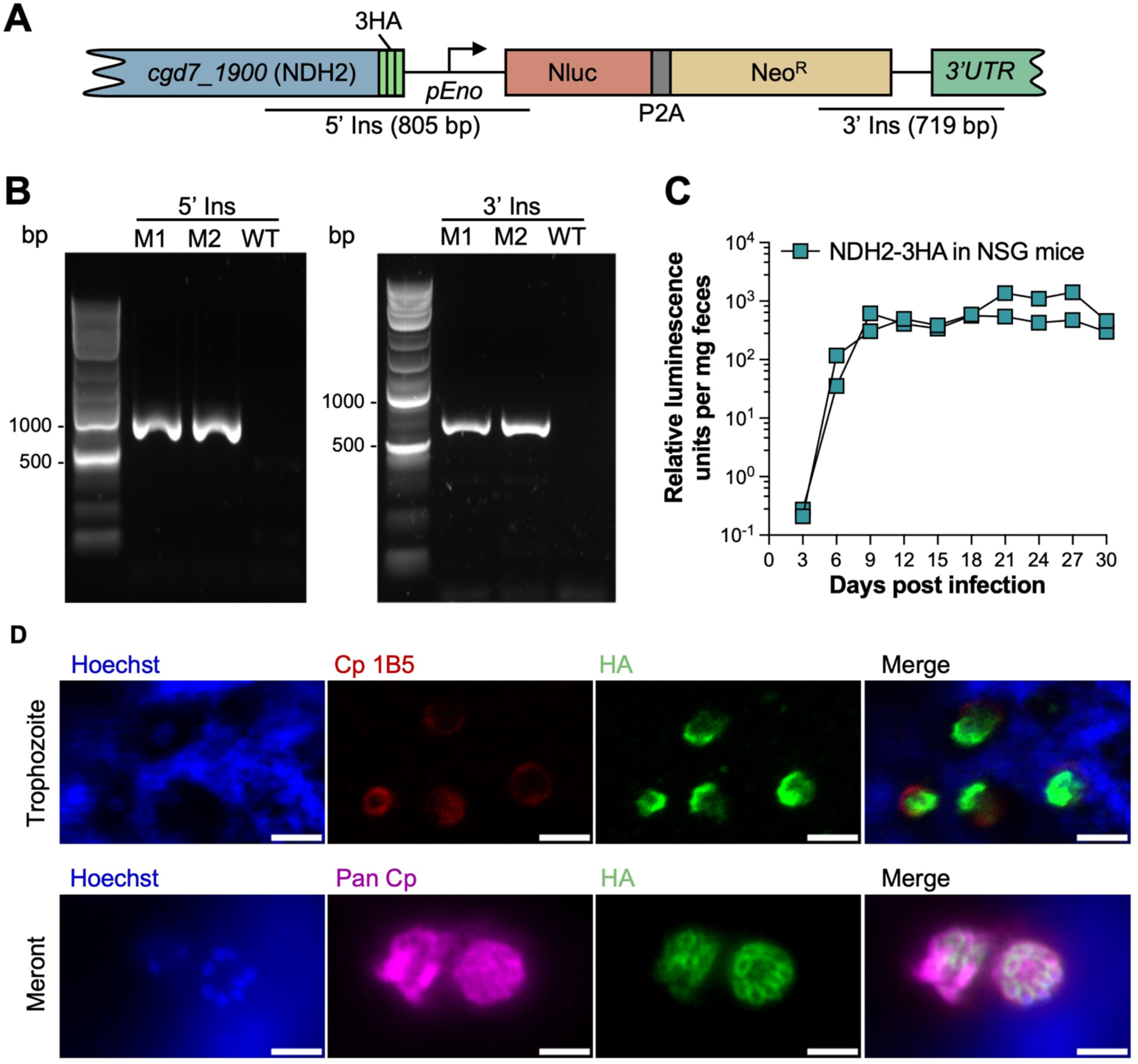
Localization of NDH2 in *C. parvum*. (A) Schematic of the NDH2-3HA-tagged endogenous locus in stable transgenic parasites. *C. parvum* sporozoites were co-transfected with NDH2-3HA-Nluc-P2A-Neo^R^ tagging plasmid and CRISPR/Cas9 plasmid containing sgRNA specific to the C terminal of NDH2. Nluc, Nanoluc luciferase; P2A, split peptide; Neo^R^, neomycin resistant cassette. 5’ Ins and 3’ Ins refer to fragments used for diagnostic PCR in B. (B) Genotype analysis of NDH2-3HA-tagged *C. parvum* strain by PCR. M1 and M2, NDH2-3HA-tagged parasites from two NSG mice; WT, wild type parasite. The product 5’ Ins is specific for the 5’ CRISPR targeting site of NDH2-3HA. The product 3’ Ins is specific for the 3’ CRISPR targeting site of NDH2-3HA. Primers are defined in **Table S1**. (C) Relative luminescence per milligram of feces from NSG mice challenged by NDH2-3HA-tagged *C. parvum*. The NDH2-3HA-tagged strain was amplified in NSG mice. Each data point represents a single fecal pellet, and each connecting line represents an individual infected NSG mouse. (D) Immunofluorescence staining of transgenic NDH2-3HA-tagged parasites. HCT-8 cells were infected with NDH2-3HA oocysts. At 24 hpi, coverslips were fixed and stained with rat anti-HA followed by goat anti-rat IgG Alexa Fluor 488 (green), mouse Cp 1B5 followed by goat anti-mouse IgG Alexa Fluor 568 (red) or Pan Cp followed by goat anti-rabbit IgG Alexa Fluor 647 (magenta), and Hoechst (blue) for nuclear staining. Scale bars, 2 µm.

### CpAOX exhibits a mitosome-like localization pattern

We also generated an epitope tagged strain to localize CpAOX. Due to its relatively low expression, we tagged CpAOX with a spaghetti monster HA tag (smHA) (32), which contains 10 separate HA epitopes on a non-fluorescent GFP protein backbone (**Fig. 3A**). The genotype CpAOX-smHA tagging strain was validated using diagnostic PCR with primers specific to the modified endogenous locus (**Fig. 3B**). Amplification of the CpAOX-smHA strain in NSG mice exhibited comparable nanoluciferase levels to those of CpNDH3-3HA parasites, although the differences in epitopes limit direct comparisons of growth efficiency (**Fig. 3C and 2C**). To visualize the location of AOX, we performed IFA and observed a punctate staining pattern of the AOX protein in both the trophozoite and meront stages (**Fig. S1**). Due to the diminutive dimensions of the mitosome, we utilized ultrastructure expansion microscopy (U-ExM), which can increase the specimen size by up to 4-fold (33). U-ExM laser scanning confocal images revealed that CpAOX was expressed in both trophozoites and meront stages and localized close to parasite nucleus, which appeared elongated in trophozoites and punctate aggregated in meronts (**Fig. 3D**). This staining pattern is highly consistent with the localization of *C. parvum* mitosome observed by electron microscopy from previous studies (34).

**Figure 3.**
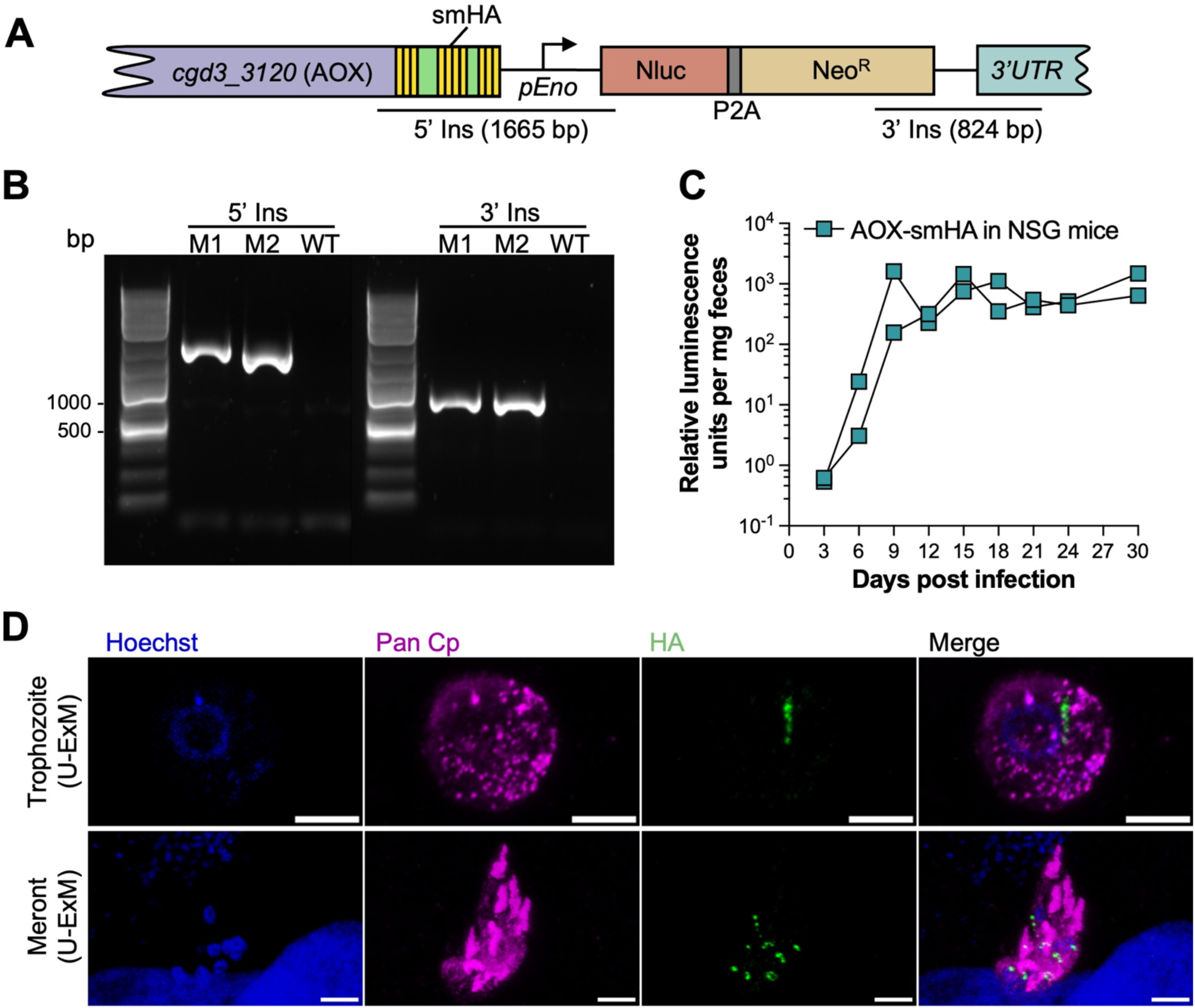
Localization of AOX in *C. parvum.* (A) Schematic of the AOX-smHA-tagged endogenous locus in stable transgenic parasites. *C. parvum* sporozoites were cotransfected with AOX-smHA-Nluc-P2A-Neo^R^ tagging plasmid and CRISPR/Cas9 plasmid containing sgRNA specific to the C terminal of AOX. smHA, spaghetti-monster HA. 5’ Ins and 3’ Ins refer to diagnostic PCR fragments used in B. (B) Genotype analysis of AOX-smHA-tagged *C. parvum* strain by PCR. M1 and M2, AOX-smHA-tagged parasites from two NSG mice. The product 5’ Ins is specific for the 5’ CRISPR targeting site of AOX-smHA. The product 3’ Ins is specific for the 3’ CRISPR targeting site of AOX-smHA. Primers are defined in Table S1. (C) Relative luminescence per milligram of feces from NSG mice challenged by AOX-smHA-tagged *C. parvum*. AOX-smHA-tagged strain was amplified in NSG mice. Each data point represents a single fecal pellet, and each connecting line represents an individual infected NSG mouse. (C) (D) U-ExM of transgenic AOX-smHA parasites at intracellular stages. HCT-8 cells were infected with AOX-smHA oocysts. At 24hpi, infected cells were fixed and expanded in gel for U-ExM. Expanded samples were stained with rabbit anti-HA followed by goat anti-rabbit IgG Alexa Fluor 488 (green), rabbit Pan Cp followed by goat anti-rabbit IgG Alexa Fluor 647 (magenta), and Hoechst (blue) for nuclear staining. Scale bars, 5 µm.

### Neither AOX nor NDH2 is essential for parasite growth

To test the essentiality of AOX or NDH2 for *C. parvum* growth, we generated knockout strains to deplete either AOX (**Fig. 4A**) or NDH2 (**Fig. 4C**) from parasites using CRISPR/Cas9. The targeted gene was replaced by a mCherry expression cassette driven by the *C. parvum* actin promoter, and the Nluc-P2A-Neo^R^ selection marker described above. Following selection and amplification in GKO and NSG mice, we successfully obtained both Λι*aox* and Λι*ndh2* strains. Deletion of the targeted genes and insertion of selective marker in specific genomic sites were validated using diagnostic PCR from fecal samples from NSG mice at 30 dpi (**Fig. 4B and 4D**). These PCR results confirmed the complete deletion of either AOX or NDH2 in the knockout strains. The fitness of knockout strains was similar to that of tagging strains based on comparison nanoluciferase assays, suggesting there is little or no deficiency in growth due to loss of either gene (**Fig. 4E, 4F, 2B and 3C**).

**Figure 4.**
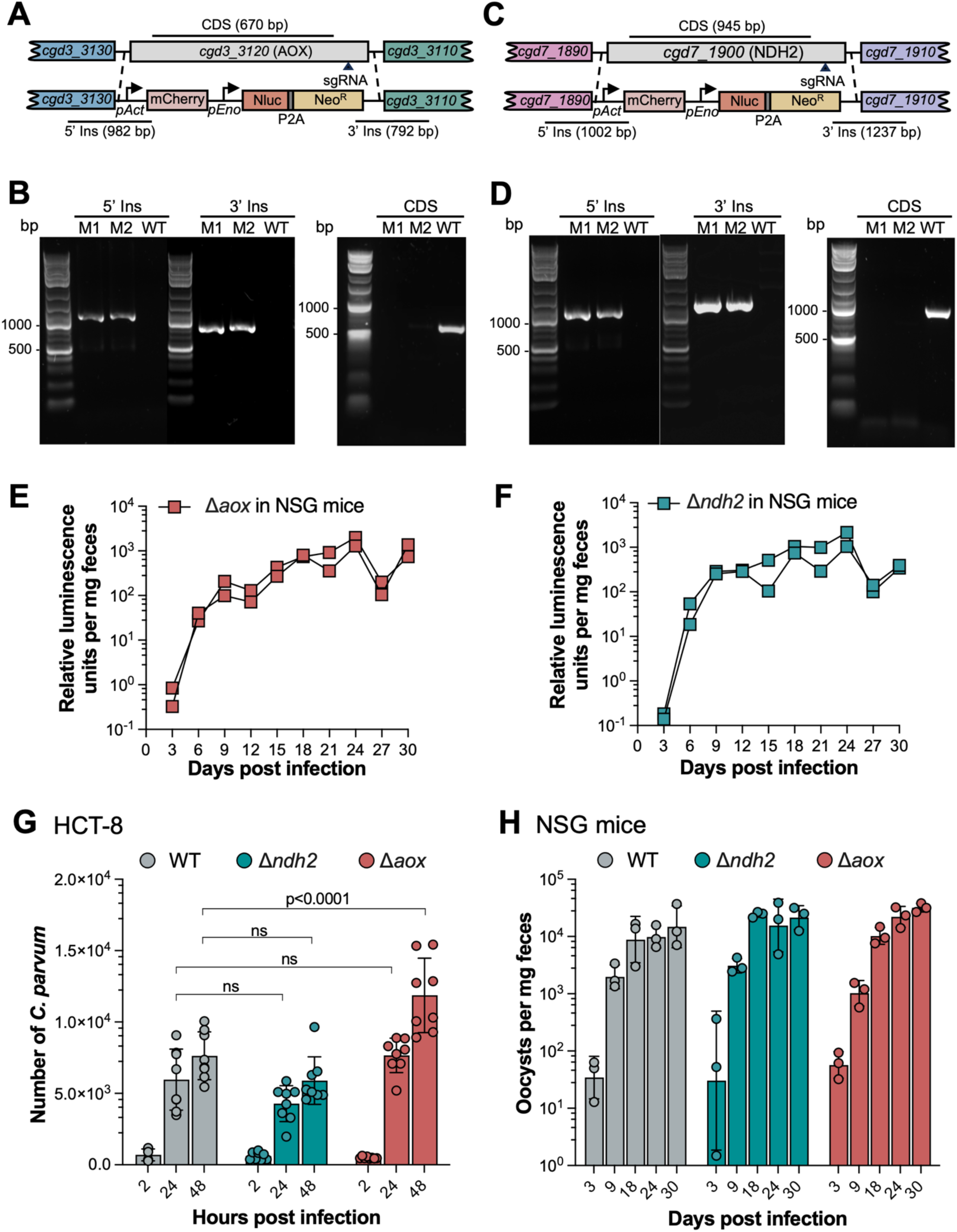
Testing essentiality of NDH2 and AOX for parasite growth. (A) Diagram of the strategy to construct Λι*aox* transgenic parasites. Construct was designed to replace the AOX locus with an mCherry and Nluc-P2A-Neo^R^ cassette. The top line shows the genomic locus and the bottom line the successfully targeted transgenic locus. sgRNA, small guide RNA. 5’ Ins and 3’ Ins refer to diagnostic PCR fragments. (B) Genotype analysis of Λι*aox C. parvum* strain by PCR. M1 and M2, Λι*aox* parasites from two NSG mice. The product 5’ Ins and 3’ Ins are specific for the 5’ CRISPR targeting site and the 3’ CRISPR targeting site for *aox* knock out, respectively. The product CDS is specific for the coding sequence of AOX. Primers are defined in **Table S1**. (C) Diagram of the strategy to construct Λι*ndh2* transgenic parasites. (D) Genotype analysis of Λι*ndh2 C. parvum* strain by PCR. M1 and M2, Λι*ndh2* parasite from two NSG mice. The product 5’ Ins and 3’ Ins are specific for the 5’ CRISPR targeting site and the 3’ CRISPR targeting site for *nd2* knock out, respectively. The product CDS is specific for the coding sequence of NDH2. Primers are defined in **Table S1**. (E) Relative luminescence per milligram of feces from NSG mice challenged by Λι*aox* parasites. Each data point represents a single fecal pellet, and each connecting line represents an individual infected NSG mouse. (F) Relative luminescence per milligram of feces from NSG mice challenged by Λι*ndh2* parasites. Each data point represents a single fecal pellet, and each connecting line represents an individual infected NSG mouse. (G) In vitro growth assay of WT, Λι*aox* and Λι*ndh2* strain. Relative fluorescence of *C. parvum* was quantified using a cell imaging reader. Values are plotted as the means ± SD. Statistical analysis was performed using two-way ANOVA with Tukey’s multi-comparison test of data from two independent experiments. ns, not significant. (H) In vivo growth assay of WT, Λι*aox* and Λι*ndh2* strain. NSG mouse was challenged with 2x10^4^ oocysts via oral gavage. 3 NSG mice was infected with each *C. parvum* strain. Fecal samples were collected at D3, D9, D18, D24, and D30 pi. DNA was extracted from fecal samples and oocyst shedding form mice was evaluated using qPCR with primers specific to *C. parvum* GAPDH. Values are plotted as the means ± SD. Statistical analysis was performed using two-way ANOVA with Tukey’s multi-comparison test of data (n=3). No statistically significant difference was detected between wild type and knock out strains at respective time points.

To further characterize the growth of knockout strains, we performed growth assays in vivo in NSG mice and in vitro with HCT-8 cells to compare the growth fitness of knockout strains to wild type parasites. To reduce the effect of host adaptation, we passaged the calve-derived wild type parasites in NSG mice and collected oocysts shed from mice. In vitro parasite growth in HCT-8 cells was determined via immunofluorescent staining followed by quantification using plate-based imaging. The result of in vitro growth assay suggested a similar growth of the Λι*ndh2* strain to the wild type strain, while the Λι*aox* strain exhibited a moderate growth enhancement at 48 hr post infection (hpi) (**Fig. 4G**). However, no difference on oocysts shedding was observed when comparing the mice challenged by either knockout strain with mice infected by wild type parasites from 3 to 30 dpi (**Fig. 4H**).

### AOX is not the drug target for SHAM and 8-HQ

Previous studies have reported that two AOX inhibitors, SHAM and 8-HQ, inhibit growth of *C. parvum* (24). This sensitivity combined with the unique presence of this enzyme in the parasite and absence in the host was used as a rationale to suggest that AOX might be a good drug target (24). To further investigate the sensitivity of SHAM-and 8-HQ-mediated inhibition, we conducted dose-response assays using drugs that were diluted in a 9-point 1:2.5 series, starting at 6 μM for SHAM and 1 μM for 8-HQ and used to treat HCT-8 cells challenged by wild type *C. parvum*. We determined EC_50_ values of SHAM and 8-HQ for *C. parvum* growth inhibition of 0.235 μM and 0.032 μM, respectively (**Fig. 5A and B**). The EC_90_ values of each drug were calculated via computational tool (https://www.graphpad.com/quickcalcs/Ecanything1/) providing estimates of 3.54 μM for SHAM and 0.12 μM for 8-HQ. To compare the sensitivity of Δ*aox* and wild type strains, we treated parasites grown in HCT-8 cells with SHAM or 8-HQ at EC_50_ or EC_90_ for 24h. The growth assay results indicated that neither of the drugs exhibited different effectivity to Δ*aox* compared with that of wild type parasites (**Fig. 5C and 5D**).

**Figure 5.**
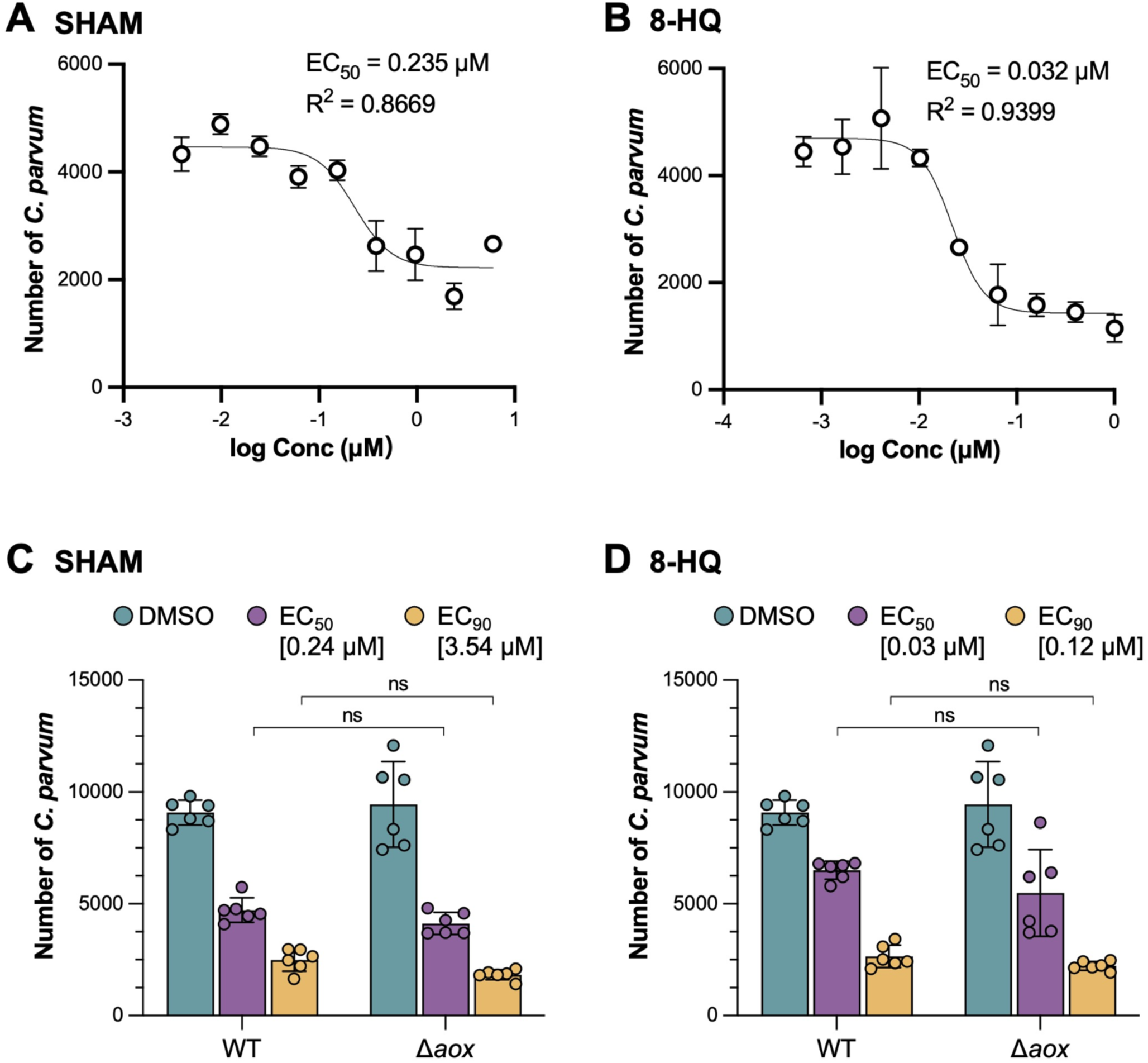
Sensitivity of Δ*aox* strain parasites to inhibitors. (A) Dose-response of *C. parvum* growth vs. concentrations of SHAM. Drugs were tested in a 9-point 1:2.5 serial dilution series starting at 6 μM. EC_50_ and R square values were calculated in GraphPad Prism 9 using a nonlinear regression curve fit. (B) Dose-response of *C. parvum* growth vs. concentrations of 8-HQ. Drugs were tested in a 9-point 1:2.5 serial dilution series starting at 1 μM. (C) Relative growth of WT and AOX-KO parasites treated with SHAM at EC_50_ or EC_90_. Plate based growth assay using HCT-8 infected cells that were fixed and stained with rat anti-HA followed by goat anti-rat IgG Alexa Fluor 488 and imaging using a Cytation 3.0 plate imager. Values are plotted as the means ± SD. Statistical analysis was performed using two-way ANOVA with Tukey’s multi-comparison test of data from two independent experiments. ns, not significant. (C) Relative growth of WT and AOX-KO parasites treated with 8-HQ at EC_50_ or EC_90_. Plate based growth assay using HCT-8 infected cells that were fixed and stained with rat anti-HA followed by goat anti-rat IgG Alexa Fluor 488 and imaging using a Cytation 3.0 plate imager. Values are plotted as the means ± SD. Statistical analysis was performed using two-way ANOVA with Tukey’s multi-comparison test of data from two independent experiments. ns, not significant.

## Discussion

*C. parvum* and *C. hominis*, the most common species infecting humans, possess a relict mitochondria-related organelle, called the mitosome, which is highly reduced in size, morphology, and functionality. Comparative genomic analysis of *C. parvum* indicates that mitochondrial metabolism-related proteins are restricted to Fe-S biosynthesis, ubiquinone biosynthesis, and an alternative ETC including MQO, NDH2, and AOX. Due to their absence in mammalian hosts, NDH2 and AOX have been proposed to be potential drug targets for cryptosporidiosis. In this study, we focused on the characterization of NDH2 and AOX and clarified that only AOX exhibited a mitosome-like localization, whereas NDH2 was found at the surface membrane or IMC. Moreover, depletion of NDH2 or AOX showed a minor effect on growth in vitro and no impact on the growth fitness of *C. parvum* in mice. Furthermore, although AOX inhibitors SHAM and 8-HQ have been reported to suppress the growth of *C. parvum* in vitro, our finding that Δ*aox* parasites are similarly sensitive to these inhibitors rules out AOX as the target of these compounds. Collectively, these findings force a revision to the proposed model for how the mitosome membrane potential is generated and also deprioritize NDH2 and AOX as potential drug targets.

Type II NDH enzymes sit in the inner leaflet of the inner mitochondrial membrane and have been reported in mitochondria of plants, fungi, as well as protists, but not in mammals (13, 35). Given its essentiality in respiration metabolism in bacterial pathogens and absence in mammalian hosts, these enzymes have been proposed as potential novel therapeutic targets (36-39). NDH2 proteins are also predicted to be encoded in the genome of apicomplexans, which lack a canonical complex I (8). Two isoforms of NDH2 are found in *T. gondii*, both of which are internal, monomeric proteins facing with their active sites to the mitochondria matrix (14). Functional analysis suggested that these two isoforms are individually non-essential; however, depletion of either isoform decreased the growth rate and reduced the mitochondrial membrane potential in *T. gondii* (14). *Plasmodium* spp. express a single NDH2 protein, which was initially reported to be sensitive to diphenylene iodonium chloride (DIC) that depolarizws the mitochondrial membrane potential leading to parasite death (18). However, this finding was challenged by later study using recombinantly expressed PfNDH2, which found that it is not sensitive to DIC (40). Moreover, a recent study depleted PfNDH2 using CRISPR/Cas9 and demonstrated that this protein is dispensable in *P. falciparum*, and that mutant is not sensitive to inhibitors of the ETC (41).

In the present study, we were surprised to discover that NDH2 in *C. parvum* is predominantly expressed at the parasite membrane. Due to limitations in the resolution of light microscopy, we are unable to differentiate between a surface membrane localization and localization in the IMC, as reported by the HyperLOPIT study (31). Given the putative enzymatic activity of NDH2, the conversion of NADH to NAD^+^ and H^+^, it is unclear why this activity would be required at the parasite surface and what reductive electron acceptor would be involved at this interface. Regardless or its exact function, depletion of this protein did not significantly affect parasite growth, either in vitro or in vivo, indicating that NDH2 is dispensable for *C. parvum* growth. This finding is consistent with the recent finding on PfNDH2 demonstrating that this protein is non-essential for *P. falciparum* growth in red blood cells (41).

Similar to NDH2, AOX has only been identified in non-mammal organisms, nominating it a potential drug target. Most plants and fungi contain both a canonical respiration pathway using complex III and IV, and an alternative respiration pathway involving AOX. In some fungi and plants, AOX genes are constitutively transcribed at a low basal level without the detectable protein and enzyme activity, whereas its expression can be activated upon the inhibition of canonical respiration pathway or presence of oxidative stress in these organisms (42, 43). Unlike NDH2, an AOX-like protein was not identified in the genomes of either *T. gondii* or *P. falciparum.* The most extensively studied AOX expressed in protozoan parasite is the *Trypanosoma brucei* AOX (TAO), which demonstrates a developmentally regulated expression in the *T. brucei* life cycle (23). As the only terminal oxidase of the mitochondrial ETC in bloodstream *T. brucei*, AOX exhibits a significant higher mRNA level and stability as well as protein abundance, compared to the procyclic form (44). Reduction in TAO mRNA level using RNAi or treatment with AOX inhibitor SHAM inhibits *T. brucei* growth (45). Similarly, AOX is also the only identified terminal oxidase in parasite genome in *C. parvum* and previous studies have suggested the sensitivity of *C. parvum* to AOX inhibitors make this a potential target for development of therapeutics (24). In the present study, we used U-ExM to examine the localization of AOX in *C. parvum* and observed staining patterns that are consistent with the mitosome, which is an elongate oval in trophozoite and a single small spherical body in mature merozoites. We have thus far been unable to confirm this localization by immuno-EM and there are current no verified makers for the mitosome that could be used for colocalization.

Surprisingly, depletion of AOX using CRISPR/Cas9 did not show any impact on the asexual development, although a modest growth enhancement was observed in AOX-depleted parasite during the late stages of the life cycle. However, no significant differences were observed in oocyst shedding from mice infected with Δ*aox C. parvum* compared to wild type parasites, indicating that AOX is non-essential for parasite growth. Previous studies have shown that AOX inhibitor SHAM and 8-HQ inhibit the *C. parvum* growth in vitro (24, 25), which was confirmed by the dose-response assays in this study. However, Δ*aox* parasites showed sensitivities to SHAM and 8-HQ that were similar to wild type parasites, indicating that AOX is not the primary target of SHAM and 8-HQ in *C. parvum*.

Our study revealed that NDH2 is surface membrane localized and non-essential in *C. parvum*. Although AOX exhibits a mitosome localization pattern, it is also dispensable for parasite growth and KO mutants are equally sensitive to the AOX inhibitor SHAM and 8-HQ. These findings challenge the previous proposal that AOX and NDH2 are potential drug target of future therapeutic development for cryptosporidiosis. Additionally, the finding that the proposed alternative ETC is nonessential, indicates that the membrane potential in the mitosome must be generated by an alternative means. The predicted localization of the MQO proteins in *C. parvum* in the nucleus or cytoplasmic fraction combined with the non-essentiality of AOX, reduces their potential importance in contributing to the membrane potential. Previous studies have suggested that TH, which can mediate proton pumping, maybe coupled to the alternative ETC to generate the membrane potential in *C. parvum* mitosome (8). However, the *C. parvum* TH protein encoded by *cgd8_2330* is predicted to localize in micronemes (31). It remains possible that the remaining TH protein encoded by *cgd1_990* is mitosome localized, although predictions from MitoProt do not support this localization (8). Alternatively, *C. parvum* contains an ADP/ATP carrier protein that is normally in the mitochondria and predicted to be in the mitosome (46). The absence of oxidative phosphorylation in *C. parvum* suggests that the ADP/ATP carrier protein may work in reverse to pump ATP into the mitosome, thus providing a source of energy for critical reactions such as iron sulfur cluster biosynthesis (8, 11, 29). Previously it was proposed that the import of ATP^-4^ in exchange for ADP^-3^ creates a charge asymmetry that generates a membrane potential in the *C. parvum* mitosome (47); similar to petite mutants in human cells lacking a functional ETC (48). Further studies are needed to resolve the localization and function of the ADP/ATP carrier protein in *C. parvum* and to resolve the mechanism by which the membrane potential is generated.

## Materials and Methods

### Animal studies

Animal studies using mice were approved by the Institutional Animal Studies Committee (School of Medicine, Washington University in St. Louis). *Ifng*^-/-^ mice (referred to as GKO) (002287; Jackson Laboratories), and Nod scid gamma mice (referred to as NSG) (005557; Jackson Laboratories) were bred in-house at Washington University School of Medicine and were separated by sex after weaning. Mice were reared in a specific-pathogen-free facility on a 12-h:12-h light-dark cycle and water ad libitum. For selection and amplification of transgenic *C. parvum* parasites, 8-to 12-week-old male or female mice were used, and water was replaced with filtered tap water containing 16 g/liter paromomycin sulfate salt (Biosynth). During infection, animals with more than 20% body weight loss or appearing debilitated were humanely euthanized.

For monitoring parasite growth in vivo, NSG mice were challenged with 2x 10^4^ parasites by oral gavage and animals were maintained on normal feed and water. Mouse fecal pellets were collected every three days post-infection. Mice were euthanized when they lost more than 20% body weight during infection.

### HCT-8 cell culture

Human ileocecal adenocarcinoma cells (HCT-8 cells; ATCC CCL-244) were cultured in RPMI 1640 medium (ATCC modification; Gibco) supplemented with 10% fetal bovine serum. The HCT-8 cells were determined to be mycoplasma negative using the e-Myco plus kit (Intron Biotechnology).

### Parasite preparation

The *C. parvum* isolate (AUCP-1) was maintained by repeated passage in male Holstein calves and purified from fecal material, as described previously (49). Purified oocysts were stored at 4°C in 50 mM Tris-10 mM EDTA (pH 7.2) for up to 6 months before use. Oocysts were prepared with 40% bleach before infection, as described previously (Xu et al., 2019). Briefly, purified oocysts were incubated with 40% bleach in DPBS (Corning Cellgro) for 10 min on ice. Oocysts were then washed 4 times in DPBS containing 1% (wt/vol) bovine serum albumin (BSA; Sigma) and resuspended in 1% BSA/DPBS. For some experiments, oocysts were excysted prior to infection by incubating the oocysts with 0.75% (wt/vol) sodium taurocholate (Sigma) at 37°C for 60 min.

Transgenic parasite was purified from NSG mice feces using saturated sodium chloride (NaCl) flotation as described in (50). Briefly, fecal pellets from infected mice were mixed and washed in cold distill water followed by centrifugation at 2,000x g for 10 min at 4°C. The pellet was resuspended in cold distill water and mixed with flotation medium (saturated NaCl solution, d = 1.18 g/ml, supplemented with 0.2% Tween-20). Cold distilled water was overlaid to prevent destruction of oocysts resulting from extended exposition to the hypertonic NaCl solution. The tube was centrifuged at 2,000x g for 30 min at 4°C. Oocysts accumulated in a white thin layer at the basis of the distilled water phase. After collection, oocysts were washed three times with 1x PBS followed by centrifugation at 2,000 × g for 10 min at 4°C. The resulting pellet contained the accumulated oocysts which were resuspended in 1x PBS and quantified using C-Chip hemocytometer (INCYTO).

### CRISPR/Cas9

To generate tagging plasmids, a 5’ homology region from the C terminus of the protein (397 bp) containing a protospacer adjacent motif (PAM) and a 3’ homology region from the 3’ UTR of the gene (400 bp) were amplified from *C. parvum* genome DNA by PCR. The triple hemagglutinin (3HA) and spaghetti monster HA (smHA) epitope tags were amplified from pCpGT1-3HA and pCpGT2-smHA, respectively (33). The previously described Nluc-P2A-neoR reporter and the pUC19 backbone was amplified from pCpGT1-3HA plasmid (33). The tagging plasmids were generated by Gibson assembly (New England BioLabs) of components described above. PAM sites were mutated by PCR amplification using primers to edit the sequence followed by treatment with KLD enzyme kit (New England BioLabs).

To generate repairing templates for gene deletions, homology repair fragments flanking the mCherry-Nluc-P2A-Neo^R^ cassette with 50 bp 5’UTR and 3’UTR homology regions for the genes of interest were PCR amplified from pINS1-mCherry-Nluc-P2A-neo-INS1 (51) with primers containing appropriate gene-specific homology regions.

To generate the CRISPR/Cas9 plasmid, a single guide RNA (sgRNA) targeting 3’ end of target genes was designed using the eukaryotic pathogen CRISPR guide RNA/DNA design tool (http://grna.ctegd.uga.edu). The pCRISPR/Cas9 backbone was amplified from previously described pACT1:Cas9-GFP, U6:sgINS1 (51). pCRISPR/Cas9-sgRNA plasmids were generated with the designed sgRNA and pCRISPR/Cas9 backbone, using Q5 site-directed mutagenesis (New England Biolabs). The same pCRISPR/Cas9-sgRNA plasmid was used for tagging and depletion of the gene of interest.

All the primers used for fragment amplifications were listed in **Table S1**. All the plasmid generated in this study were described in **Table S2**.

For transfection, oocysts (1.25x 10^7^ per transfection) were excysted as described above, and sporozoites were collected by centrifugation at 2,500 rpm for 3 min and resuspended in SF buffer (Lonza) containing 50 mg of tagging plasmid or 30 mg of linear targeting template and 30 mg CRISPR/Cas9 plasmid in a total volume of 100 ml. The mixtures were then transferred to a 100 ml cuvette (Lonza) and electroporated on an AMAXA 4D-Nucleofector system (Lonza) using program EH100. Electroporated sporozoites were transferred to cold DPBS and kept on ice before infecting mice. All the repairing templates and CRISPR/Cas9 plasmids used for transgenic strains were specified in Table S3.

### Selection and amplification of transgenic parasites in mice

GKO mice were used for the first round of transgenic parasite selection. Each mouse was orally gavaged with 200 μL of 8% (w/v) sodium bicarbonate 5 min prior to infection. Mice were then gavaged with 2.5x 10^7^ electroporated sporozoites. All mice received drinking water with 16 g/L paromomycin continuously from the 1 dpi, based on previously published protocol (Vinayak et al., 2015). Fecal pellets were collected begin at 9 to 15 dpi, after which animals were euthanized by CO_2_ asphyxiation according to the animal protocol guidelines. Fecal pellets were stored at -80°C for qPCR or at 4°C for luciferase assays or for isolating oocysts for subsequent infections. A second round of amplification was performed by orally gavaging NSG mice using a fecal slurry from GKO mice described above. The fecal pellets were transferred to a 1.7 mL microcentrifuge tube, ground with a pestle, diluted by addition of 1 mL cold 1x DPBS, vortexed for 30 s followed by a centrifugation at 200 rpm for 10 min to pellet large particulates. Oocysts in the supernatant was counted using C-Chip hemocytometer and diluted in 1x DPBS. 2x 10^4^ oocysts were gavaged into one NSG mouse. Infected NSG mice were treated with 16 g/L paromomycin drinking water for the entirety of the experiment. Fecal pellets for qPCR and luciferase assay were collected every 3 days starting 3 dpi and fecal pellets for purification were collected every day starting at 12 dpi and stored at 4°C. Oocyst purification from NSG feces was as described above. Purified oocysts were stored in PBS at 4°C and used within 6 months of extraction.

### Luciferase assay

Luciferase assays were performed using the Nano-Glo Luciferase assay kit (Promega). Mouse fecal pellets were collected and weight in 1.7-ml microcentrifuge tubes, ground with a pestle. Glass beads (3 mm; Fisher Scientific) and 1 ml fecal lysis buffer (50 mM Tris pH 7.6, 2 mM DTT, 2 mM EDTA pH 8.0, 10% glycerol, 1% Triton X-100 prepared in water) (Pawlowic et al., 2017) were added to the tube for fecal sample lysis. Tubes were incubated at 4°C for 30 min, vortexed for 1 min, and then spun at 16,000x g for 1 min to pellet debris. 100 mL supernatant was added to one well of a 96-well white plate (Costar 3610) with two technique replicates for each sample, and then 100 mL of a 25:1 Nano-Glo Luciferase buffer to Nano-Glo Luciferase substrate mix was added to each well. The plate was incubated in dark for 3 min at room temperature. Luminescence values were read on a Cytation 3 cell imaging multi-mode reader (BioTek).

### Fecal DNA extraction and quantification of oocysts using qPCR

DNA was extracted from fecal pellets using the QIAamp PowerFecal DNA kit (Qiagen) according to the manufacturer’s protocol. Oocyst numbers were quantified using qPCR with the *C. parvum* glyceraldehyde-3-phosphate dehydrogenase (GAPDH) primers (**Table S1**), as described previously (Wilke et al., 2019). A standard curve was established by purifying genomic DNA from a known number of oocysts following a serial dilution. Reactions were performed on a QuantStudio 3 real-time PCR system (Thermo Fisher) with the amplification conditions as previously described (Wilke et al., 2019).

### Genotyping of transgenic parasites

To check for the successful insertion of the target sequence into the genomic locus of specific gene, PCR was performed on 1ml purified fecal DNA using Q5 Hot Start high-fidelity 2x master mix (New England Biolabs) with primers listed in **Table S1**. PCRs were performed on a Veriti 96-well thermal cycler (Applied Biosystems) with the following cycling conditions: 98°C for 30 s, followed by 35 cycles of 98°C for 15 s, 60°C for 30 s, and 72°C for 2 min, with a final extension of 72°C for 2 min. Melting temperature and extension time may vary from different PCR reaction for the specific primers and distinct product length. PCR products were resolved on 1.0% agarose gel containing GelRed (diluted 1:10,000; Biotium) and imaged on a ChemiDoc MP imaging system (Bio-Rad).

### Indirect immunofluorescence microscopy

HCT-8 cells grown on coverslips with 80% confluency were infected with 1x 10^5^ oocysts per well. At specific time points postinfection, infected cells were fixed with 4% formaldehyde for 10 min and washed three times with PBS. The fixed samples were then permeabilized and blocked with blocking buffer consisting of 1% BSA and 0.1% Triton X-100 (Sigma) in PBS. Primary antibodies were diluted in blocking buffer: rat anti-HA was used at 1:500, rabbit anti-HA was used at 1:500 (for U-ExM), MAb 1B5 (hybridoma supernatant) was used at 1:250, and Pan Cp (rabbit polyclonal antibody) was used at 1:10,000. Cells were incubated with primary antibodies for 1 h at room temperature, washed three times with PBS, and then incubated for 1 h at room temperature in secondary antibodies conjugated to Alexa Fluor dyes (Thermo Fisher Scientific) diluted 1:1,000 in blocking buffer. Nuclear DNA was stained with Hoechst (Thermo Fisher Scientific) diluted 1:1,000 in blocking buffer for 20 min at room temperature and then mounted with Prolong Glass antifade mountant (Thermo Fisher Scientific). Images were captured on a Zeiss Axioskop Mot Plus fluorescence microscope equipped with a 100x, 1.4 N.A. Zeiss Plan Apochromat oil objective lens or on a Zeiss LSM880 laser scanning confocal microscope equipped with a 63x, 1.4 N.A. Zeiss Plan Apochromat oil objective lens. Images were acquired using AxioVision Rel v 4.8, software or ZEN v2.1, v2.5 software. Images were adjusted in ImageJ v2.0.0 (https://fiji.sc/).

### Expansion microscopy

U-ExM was applied as described previously (52). HCT-8 cells were infected and fixed the same way as described for immunofluorescence staining. Samples were embedded overnight in a mixture of 1% acrylamide and 0.7% formaldehyde for protein anchor and crosslinking prevention. The formed gel was transferred into denaturation buffer and denatured at 95°C. Polymerization of expansion gel was performed on ice containing monomer solution (19% sodium acrylate/10% acrylamide/0.1% (1,2-Dihydroxyethylene) bisacrylamide), 0.5% ammonium persulfate (APS) and 0.5% tetramethyl ethylenediamine (TEMED). Polymerized gels were denatured at 95°C for 90 min in the denaturation buffer (200 mM SDS, 200 mM NaCl, 50 mM Tris-Base, pH= 9.0) and expanded in pure H_2_O overnight. On the next day, the expansion ratio of fully expanded gels was determined by measuring the diameter of gels. Well expanded gels were shrunk in PBS and stained with primary (rabbit anti-HA at 1:200, rat Pan Cp at 1:500), secondary antibodies (Alexa Fluor dyes at 1:500), and Hoechst (at 1:500) diluted in freshly prepared PBS/BSA 2% at room temperature for 6 h. Three washes with PBS/0.1% Tween for 10 min were performed after each staining. Stained gels were expanded again in pure H2O overnight for further imaging. Images were captured on a Zeiss LSM880 laser scanning confocal microscope equipped with a 63x, 1.4 N.A. Zeiss Plan Apochromat oil objective lens and acquired using ZEN v2.1, v2.5 software.

### *C. parvum* growth assay and drug treatment in vitro

HCT-8 cells were plated at 1 x 10^5^ cells per well in black-sided, optically clear-bottomed 96-well plates (Greiner Bio-One) and grown for 24 h until confluent. Cells were infected with 5 x 10^3^ bleached oocysts per well. After 24 h of infection/treatment, cells were fixed in 4% formaldehyde for 10 min, washed three times with PBS, and then permeabilized and blocked in PBS containing 0.1% Triton X-100 and 1% BSA for 20 min. *C. parvum* parasites were labeled with rabbit Pan Cp diluted 1:2,000 in blocking buffer followed by Alexa Fluor goat anti-rabbit 488 secondary antibody (1:1000). Host cell nuclei were stained with Hoechst for 20 min. Plates were imaged with a 10x objective on a BioTek Cytation 3 cell imager (20 images per well in a 5 x 4 grid). BioTek Gen5 software version 3.08 (Agilent) was used to quantify the total number of parasites (puncta in the GFP channel) and host cells (nuclei in the DAPI channel) per well.

For dose-response *C. parvum* growth inhibition assay of SHAM (ThermoFisher) and 8-HQ (ThermoFisher) for *C. parvum*, drugs were tested in a 9-point 1:2.5 serial dilution series starting at 6 uM (SHAM) and 1 uM (8-HQ). *C. parvum* growth assays were performed as described above. EC_50_ and EC_90_ values were calculated in GraphPad Prism 9 using a nonlinear regression curve fit with six replicates per data point (three technical replicates from two independent experiments).

### Statistical analysis

All statistical analyses were performed in GraphPad Prism 10 (GraphPad Software) unless otherwise specified. Two-way ANOVA with Tukey’s multi-comparison test was performed for statistical analysis based on data from at least two biological replicates. *P* values of ≤ 0.05 were considered statistically significant.

## Data availability

All of the data are found in the manuscript or supplemental material.

## Supplemental Material

**Figure S1.**
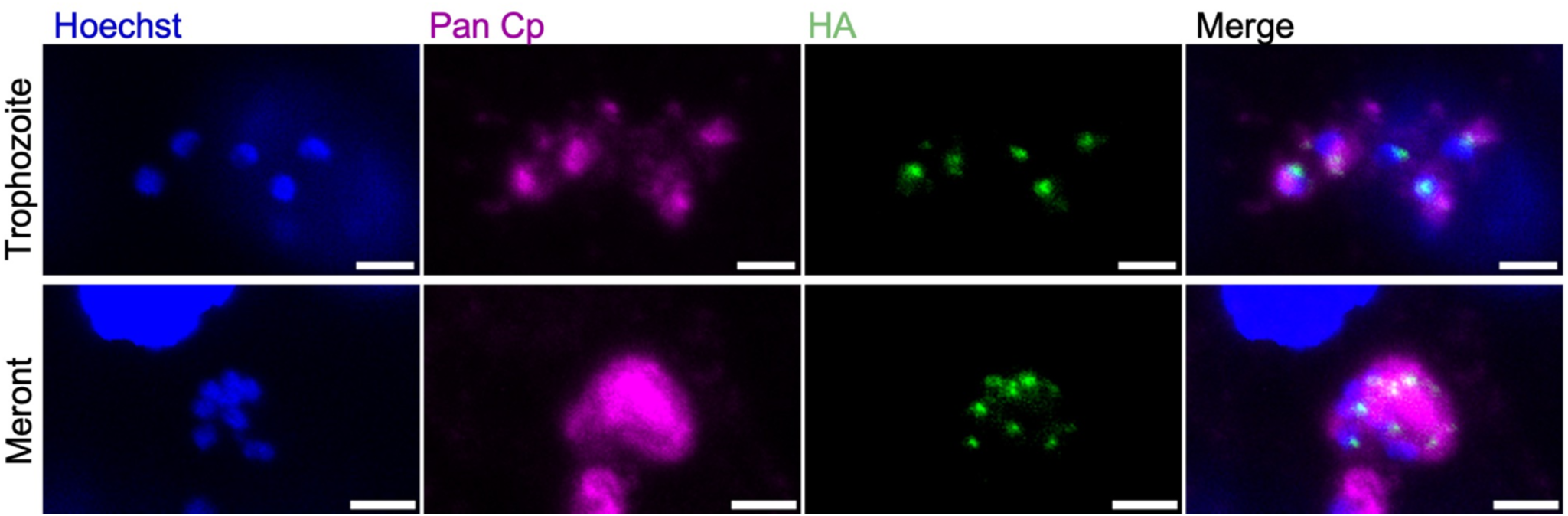
Localization of AOX in *C. parvum* using IFA. Immunofluorescence staining of transgenic AOX-smHA-tagged parasites. HCT-8 cells were infected with AOX-smHA oocysts. At 24 hpi, coverslips were fixed and stained with rat anti-HA followed by goat anti-rat IgG Alexa Fluor 488 (green), Pan Cp followed by goat anti-rabbit IgG Alexa Fluor 647 (magenta), and Hoechst (blue) for nuclear staining. Scale bars, 2 µm.

**Table S1** Primers used in the present study.

**Table S2** Plasmids used in the present study.

**Table S3** Transgenic *C. parvum* strains.

## Acknowledgements

We thank members of the Sibley laboratory for helpful comments. We are grateful to Dr. William Witola for supply oocysts of *C. parvum*. This work was supported by a grant from the National Institutes of Health NIAID (AI145496).

