## Supplemental Tables for "Function of the alternative electron transport chain in the *Cryptosporidium parvum* mitosome"

**Table S1** Primers used in the present study

| Fragment | Template | Forward primer (5' → 3') | Reverse primer (5' → 3') |
|---|---|---|---|
| EnoP-Nluc-P2A-Neo$^{R}$ | pCpGT1-3HA | ATGCATCTTCATTTAGTATCTTAGGTCG | AATTAAGATAAAAAGAAAAACTTAATCGATAC |
| pUC19 backbone | pCpGT1-3HA | GGGGATCCTCTAGAGTCGAC | GGGTACCGAGCTCGAATT |
| smHA cassette / 3HA cassette | pCpGT1-3HA/ pCpGT2-smHA | GCTAGCAAGGGCTCGGGC | GGCGCGCCAAATAAAGTAAAGTTTATCG |
| pCRISPR/Cas9 backbone | pCpGT1-3HA | GTTTTAGAGCTAGAAATAGCAAG | CCCAACACTTAACCTTTCAGT |
| AOX-5'homology arm | *C. parvum* genomic DNA | GAATTCGAGCTCGGTACCCTTACTCTACTTGAAGAGGCTGAA | GAGCCCGAGCCCTTGCTAGCTTCTCCATTTAATCTTATATCTGCG |
| AOX-3'homology arm | *C. parvum* genomic DNA | TTTTTCTTTTTATCTTAATTTAATTTAGATATTTATTGAAATATTTTATTTCATTC | GTCGACTCTAGAGGATCCCCGTCTCATCATCAATTACAACAAG |
| AOX-PAM mutation | - | AGAGAAAAGCtccGAAATTTGC | CAAATCCTGGAAGAAGACC |
| AOX-sgRNA1-Linker | - | ctgaaaggttaagtgttgggAGTAGACGGAGGCAAATTTCgttttagagctagaaatagc | |
| NDH2-5'homology arm | *C. parvum* genomic DNA | TGAATTCGAGCTCGGTACCCTCTCACGGAAGAAGACTTC | GAGCCCGAGCCCTTGCTAGCGTGAGAAACGTTCATTTTGTAG |
| NDH2-3'homology arm | *C. parvum* genomic DNA | TTTTTCTTTTTATCTTAATTAAAGAAGAGATTTGACTATTTTTTG | GTCGACTCTAGAGGATCCCCCTGTTAATGGACTTTTGGC |
| NDH2-PAM mutation | - | CTACTTTGAAtccAAGTTCAAG | TATATGCTAATCGCCAGATATAC |
| NDH2-sgRNA1-linker | - | ctgaaaggttaagtgttgggAACTGCGGGATCTTGAACTTgttttagagctagaaatagc | |
| AOX 5'UTR-mCherry-Nluc-P2A-Neo$^{R}$-AOX 3'UTR | pINS1-mCherry-Nluc-P2A-neo-INS1 | TACAGCGGAATCTTATTTACAATCGTATTTTTTTTTAAATAATATTAATgctcagaatgagttggttataaac | AACTAAATTATTTTGAATGAAATAAAATATTTCAATAAATATCTAAATTAgcttaattaatcagaagaattcgtc |
| NDH2 5'UTR-mCherry-Nluc-P2A-Neo$^{R}$-NDH2 3'UTR | pINS1-mCherry-Nluc-P2A-neo-INS1 | ATTAGAAATTAATTTCAAAAAAGAAATTCTTAGTAAATAAACTTTAAATTgctcagaatgagttggttataaac | GGCTAAATAAGATGAATTGTACACACAAAAAATAGTCAAATCTCTTCTTTgcttaattaatcagaagaattcgtc |

**Genotyping of transgenic strains with NSG mice fecal DNA**

| | Forward primer (5' → 3') | Reverse primer (5' → 3') |
|---|---|---|
| AOX-smHA 5'Ins | GGTTGGTGCAATGCTTAGAC | GTGTGTGTGAAAAGCTGTC |
| AOX-smHA 3'Ins | GCTGAAGAACTTGGTGGTGA | GCCAATGCCGTCGTAAATAG |
| NDH2-3HA 5'Ins | GTTAGACCACGAAAGTTAGCAG | TGTGTGTGAAAAGCTGTC |
| NDH2-3HA 3'Ins | GCTGAAGAACTTGGTGGTGA | ACTATTGATCCAATGACTCTCTC |

| | | |
|---|---|---|
| AOX-KO 5'Ins | CCAGCCATTGGAATTATGTCC | GTCATTTTTATAGCCCTAACGC |
| AOX-KO 3'Ins | GCTGAAGAACTTGGTGGTGA | GCTTATTATTACTTCGCCAATGC |
| AOX-KO CDS | AGGCATCTTTGGCTACTCTCT | AGAAGGCAAACTGAGTCCCA |
| NDH2-KO 5'Ins | CTGCTGGAGTTTCCTCAATTAC | GTCATTTTTATAGCCCTAACGC |
| NDH2-KO 3'Ins | GCTGAAGAACTTGGTGGTGA | CGTAGAGTCAACAGTCAATGG |
| NDH2-KO CDS | GACGGCCAAAGGTTCTCATCT | CGCAATCTCCAAGTGCGTATG |
| **Oocyst quantification** | | |
| | Forward primer (5' → 3') | Reverse primer (5' → 3') |
| CpGAPDH | GAAGATGCGCTGGGAACAAC | CGGATGGCCATACCTGTGAG |

**Table S2** Plasmids used in the present study.

| Plasmid Full Name | Fragments used | Usage |
|---|---|---|
| pNDH2-3HA-Nluc-P2A-$Neo^R$ | pUC19 backbone | NDH2-3HA-tagging repairing plasmid |
| | NDH2-5'homology arm | |
| | 3HA cassette | |
| | EnoP-Nluc-P2A-$Neo^R$ | |
| | NDH2-3'homology arm | |
| pAOX-smHA-Nluc-P2A-$Neo^R$ | pUC19 backbone | AOX-smHA-tagging repairing plasmid |
| | AOX-5'homology arm | |
| | smHA cassette | |
| | EnoP-Nluc-P2A-$Neo^R$ | |
| | NDH2-3'homology arm | |
| AOX: mCherry-Nluc-P2A-$Neo^R$-AOX | AOX 5'UTR-mCherry-Nluc-P2A-$Neo^R$-AOX 3'UTR | AOX-Knockout repairing template (PCR product) |
| NDH2: mCherry-Nluc-P2A-$Neo^R$-NDH2 | NDH2 5'UTR-mCherry-Nluc-P2A-$Neo^R$-NDH2 3'UTR | NDH2-Knockout repairing template (PCR product) |
| pCRISPR/Cas9-AOX-sgRNA1 | pCRISPR/Cas9 backbone | CRISPR/Cas9 plasmid containing sgRNA specific to AOX |
| | AOX-sgRNA1-Linker | |
| pCRISPR/Cas9-NDH2-sgRNA1 | pCRISPR/Cas9 backbone | CRISPR/Cas9 plasmid containing sgRNA specific to NDH2 |
| | NDH2-sgRNA1-Linker | |

**Table S3** Transgenic *C. parvum* strains

| Name | Plasmids used for co-transfection | Usage |
|---|---|---|
| NDH2-3HA | pNDH2-3HA-Nluc-P2A-Neo$^{R}$ | NDH2-3HA-tagging |
| | pCRISPR/Cas9-NDH2-sgRNA1 | |
| AOX-smHA | pAOX-smHA-Nluc-P2A-Neo$^{R}$ | AOX-smHA-tagging |
| | pCRISPR/Cas9-AOX-sgRNA1 | |
| $\Delta$*aox* | AOX-mCherry-Nluc-P2A-Neo$^{R}$-AOX | *aox* knockout |
| | pCRISPR/Cas9-AOX-sgRNA1 | |
| $\Delta$*ndh2* | NDH2-mCherry-Nluc-P2A-Neo$^{R}$-NDH2 | *Ndh2* knockout |
| | pCRISPR/Cas9-NDH2-sgRNA1 | |
